# High prevalence and diversity of Extended-Spectrum β-Lactamase and emergence of Carbapenemase producing *Enterobacteriaceae* spp in wildlife in Catalonia

**DOI:** 10.1101/510123

**Authors:** Laila Darwich, Anna Vidal, Chiara Seminati, Andreu Albamonte, Alba Casado, Ferrán López, Rafael A. Molina-López, Lourdes Migura-Garcia

## Abstract

In wildlife, most of the studies focused on antimicrobial resistance (AMR) describe *Escherichia coli* as the principal indicator of the selective pressure. In the present study, new species of *Enterobacteriaceae* with a large panel of cephalosporin resistant (CR) genes have been isolated from wildlife in Catalonia. A total of 307 wild animals were examined to determine CR enterobacteria prevalence, AMR phenotypes and common carbapenem and CR gene expression. The overall prevalence of CR-phenotype was 13% (40/307): 17.3% in wild mammals (18/104) and 11.5% in wild birds (22/191) (p<0.01)). Hedgehogs presented the largest prevalence with 13.5% (14/104) of the mammal specimens, followed by raptors with 7.3% (14/191) of the total bird specimens. Although CR *E. coli* was obtained most frequently (45%), other CR-*Enterobacteriaceae* spp like *Klebsiella pneumoniae* (20%), *Citrobacter freundii* (15%), *Enterobacter cloacae* (5%), *Proteus mirabilis* (5%), *Providencia* spp (5%) and *Serratia marcescens* (2.5%) were isolated. A high diversity of CR genes was identified among the isolates, with 50% yielding *bla*CMY-2, 23% *bla*SHV-12, 20% *bla*CMY-1 and 18% *bla*CTX-M-15. Additionally, new CR-gene variants and resistance to carbapenems associated to OXA-48 were found. Most of the CR isolates, principally *K. pneumoniae* and *C. freundii*, were multiresistant with co-resistance to fluoroquinolones, tetracycline, sulphonamides and aminoglycosides. This study describes for the first time in wildlife a high prevalence of *Enterobacteriaceae* spp harbouring a large variety of carbapenem and CR genes frequently associated to nosocomial human infections. Implementation of control measures to reduce the impact of anthropogenic pressure in the environment is urgently needed.

## Introduction

In the last decades, the prevalence of opportunistic and antimicrobial resistant (AMR) bacteria associated with nosocomial infections has suffered an important increase in hospital settings. The overuse of antibiotics in human and veterinary medicine have led to the spread of AMR pathogens, becoming a global health problem [1].

Extended-spectrum β-lactamases (ESBLs) and plasmid-mediated AmpC-type β-lactamases (pAmpC) are the most common enzymes that confer resistance to broad-spectrum cephalosporins among members of the family *Enterobacteriaceae*. These β-lactamases have extensively diversified in response to the clinical use of new generation drugs: cephalosporins, carbapenems and monobactams [2]. These enzymes are mostly encoded by genes located in plasmids that can be horizontally transferred to different bacteria genera [1]. Carbapenems are last-line beta-lactam antibiotics with the broadest spectrum of activity. Unfortunately, carbapenems nowadays are commonly used in hospital settings for the treatment of life-threatening infections caused by cephalosporin resistant (CR) *Enterobacteriaceae*. However, the emergence of resistance to carbapenems mediated by the production of carbapenemases has led to limited therapeutic options in human health [3], with the OXA-48 variant being highly prevalent in human clinical infections [4].

The dissemination of CR has been studied widely in *Enterobacteriaceae* from humans and livestock, whereas studies concerning the environment, including wildlife, are still lacking [2]. In recent years, an important increase of CR *Escherichia coli* has been reported in different epidemiological settings such as humans, pets, livestock, retail meat and the environment [5-10]. The study of wildlife as sentinel of the AMR environmental contamination has recently acquired more consideration worldwide [11].

However, most of the environmental-wildlife interface studies have been focused on wild birds, as principal AMR disseminators by their migratory routes, with a limited variety of AMR bacteria species described. Isolation of CR-carrying bacteria from wild birds has been globally reported in *Escherichia coli* [12-17] and less frequently in *Klebsiella pneumoniae* [18]. All these results confirm the dissemination success of ESBL *bla*_SHV-12_ and *bla*_CTX-M_ variants in wild birds worldwide. More recently, presence of CR *E. coli* has also been described in wild mammals, but at lower prevalence in comparison with wild birds [19].

In the present study, we report for the first time in Spain, the presence of diverse families of CR- -encoding genes in a large variety of *Enterobacteriaceae* species including *E. coli, K. pneumoniae, Citrobacter freundii, Enterobacter cloacae, Serratia marcescens and Proteus mirabilis*- in wild mammals and wild birds. Furthermore, we describe the presence of carbapemenase resistant *E. coli* and *P. mirabilis* associated with the presence of OXA-48 in isolates of wildlife origin. These bacterial species are frequently associated with severe nosocomial infections in human hospitals of Catalonia [20].

## Material and Methods

### Study population

Wild animals attended at the Wildlife Rehabilitation Centre (WRC) of Torreferrusa (Catalonia, North-East Iberian Peninsula) were analysed between November 2016 and May 2017. All animals were examined and tested using cloacal or rectal swabs on arrival at the centre before receiving any pharmacologic or antimicrobial treatment. The anthropogenic origin was confirmed as the most frequent cause of hospitalization, comprising direct persecution (gunshot, poisoning, illegal captivity or traps) to involuntary human induced threats (collisions with vehicles, fences or electric lines and electrocution). The rehabilitation centre is under the direction of the Catalan Wildlife-Service, who stipulates the management protocols and Ethical Principles according to the Spanish legislation [21].

### Microbiological analysis

Rectal and cloacal swabs were plated in MacConkey agar supplemented with ceftriaxone (1mg/L). Single colonies growing on the plate were subculture and identified biochemically using the API (bioMérieux, Marcy l’Etoile, France) or the VITEK 2 (bioMérieux, Spain) systems. Serovar identification and phage typing of *Salmonella* spp. were carried out at the Spanish National Reference Laboratory (Algete, Madrid, Spain).

### Antimicrobial susceptibility testing

Minimal inhibitory concentration (MIC) was performed using a broth microdilution method (VetMIC GN-mo, SVA, Sweden) for the following antimicrobials: ampicilin (Am), cefotaxime (Ctx), ceftazidime (Caz), ciprofloxacin (CIP), nalidixic acid (NAL), gentamicin (GN), streptomycin (ST), kanamycin (KM), florfenicol (FF), chloramphenicol (CF), tetracycline (TE), colistin (COL), sulphametoxazole (SU) and trimethoprim (TM). The *E. coli* ATCC 25922 was used as the control strain. Epidemiological cut-off values were determined following the European Committee on Antimicrobial Susceptibility testing (EUCAST) recommendations. For those *Enterobacteriaceae* species with no cut-off values defined, cut-off values were obtained from the British Society for Antimicrobial Chemotherapy (BSAC) or the Société Française de Microbiologie (SFM).

### Characterization of antimicrobial resistance genes

Molecular diagnosis of CR genes was performed for the following genes; *bla*_SHV_, *bla*_CTX-M_, *bla*_CMY1_, *bla*_CMY2_, and *bla*_TEM_, [22], *bla*_OXA_ *bla*_VIM_ [23] and *mcr-*1 colistin-resistance genes [24].

PCR products were Sanger sequenced for verification at the Genomic and Bioinformatics Service of the Universitat Autònoma de Barcelona (Barcelona, Spain). Sequences and chromatograms were manually explored to trim bad-quality bases with BioEdit 7.2. Once the assembly of the consensus sequences was done, both complete and partial genomes were aligned using Clustal Omega program, and finally blasted against the public database (National Center for Biotechnology Information, NCBI).

### Statistical analysis

Descriptive analysis was performed under 95% confidence, using SPSS Advanced Models TM 15.0 (SPSS Inc. 233 South Wacker Drive, 11th Floor Chicago, IL 60.606-6412). The Chi-square test or Fisher exact test was used for comparison between proportions when appropriate. Statistically significant results were considered when P < 0.05.

## Results

The sample size comprised 307 wild animals belonging to 67 different species grouped as, birds (62%), mammals (34%) and reptiles (4%) (Fig 1). Animals came from different regions of Catalonia with a high density of urban areas and pig farming production.

**Fig 1.**
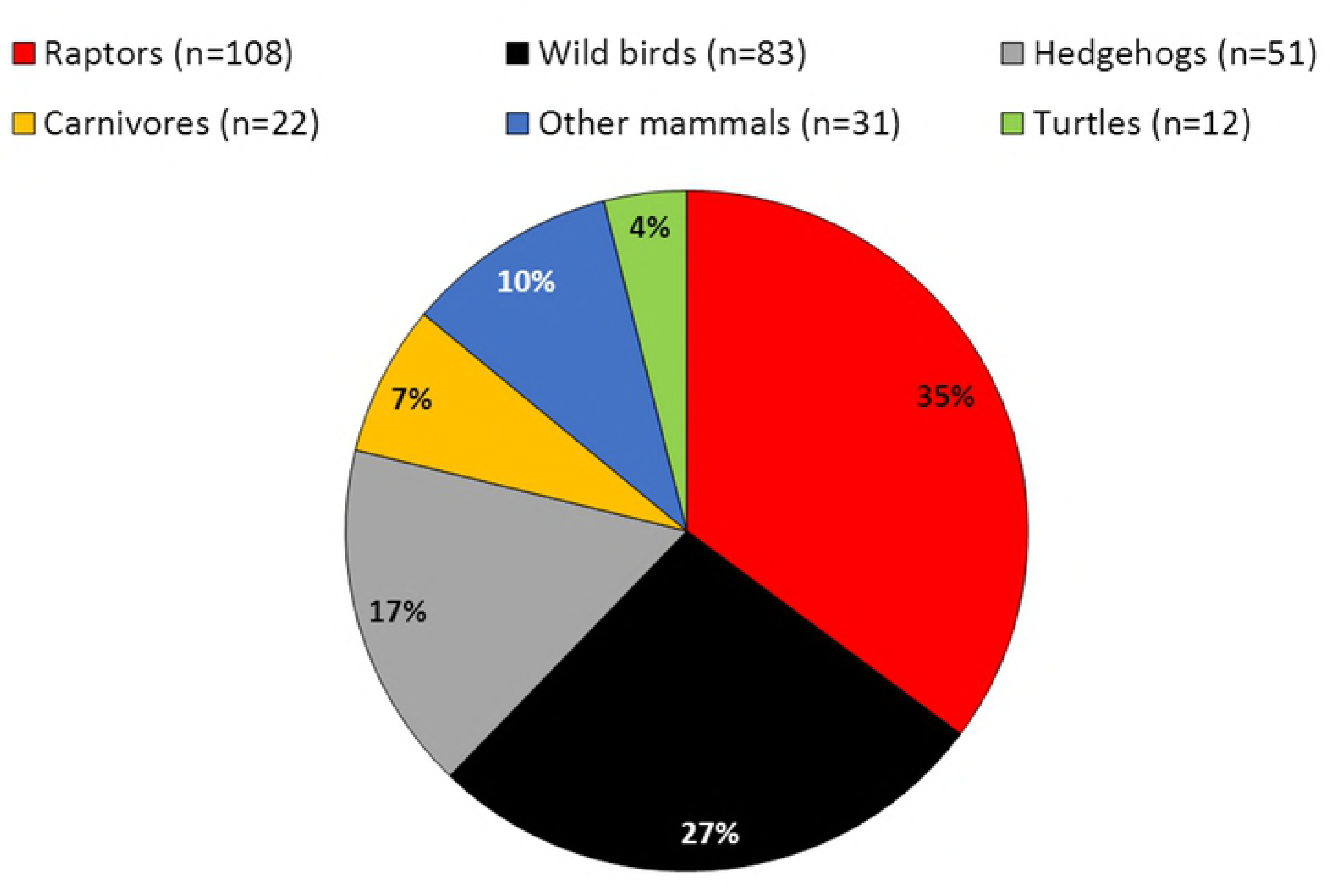
Proportion of wildlife analysed in the study according to the zoological category. Animal groups: raptors (different species of birds of prey and owls), wild birds (principally passerines and seagulls), hedgehogs (European and Algerian hedgehogs), carnivores (mainly mustelids), and other mammals (wild boars and roe deer).

Ceftriaxone resistant isolates were detected in 65 out of the 307 (21%) faecal samples analysed. Of those, 40 harboured ESBL or pAmpC-encoding genes, representing an overall prevalence of 13% (Fig 2). The prevalence of CR-carrying isolates was 17.3% in wild mammals (18/104) and 11.5% in wild birds (22/191) (p<0.01)). Surprisingly, hedgehogs presented the largest prevalence with 13.5% (14/104) of the mammal specimens [67% of the Algerian (2/3) and 26% of the European (12/47) samples harbouring CR-genes]. Within the bird group, raptors represented the highest prevalence with 7.3% (14/191) of the total bird specimens [13% (14/108) of the raptor species examined] (Fig 2).

**Fig 2.**
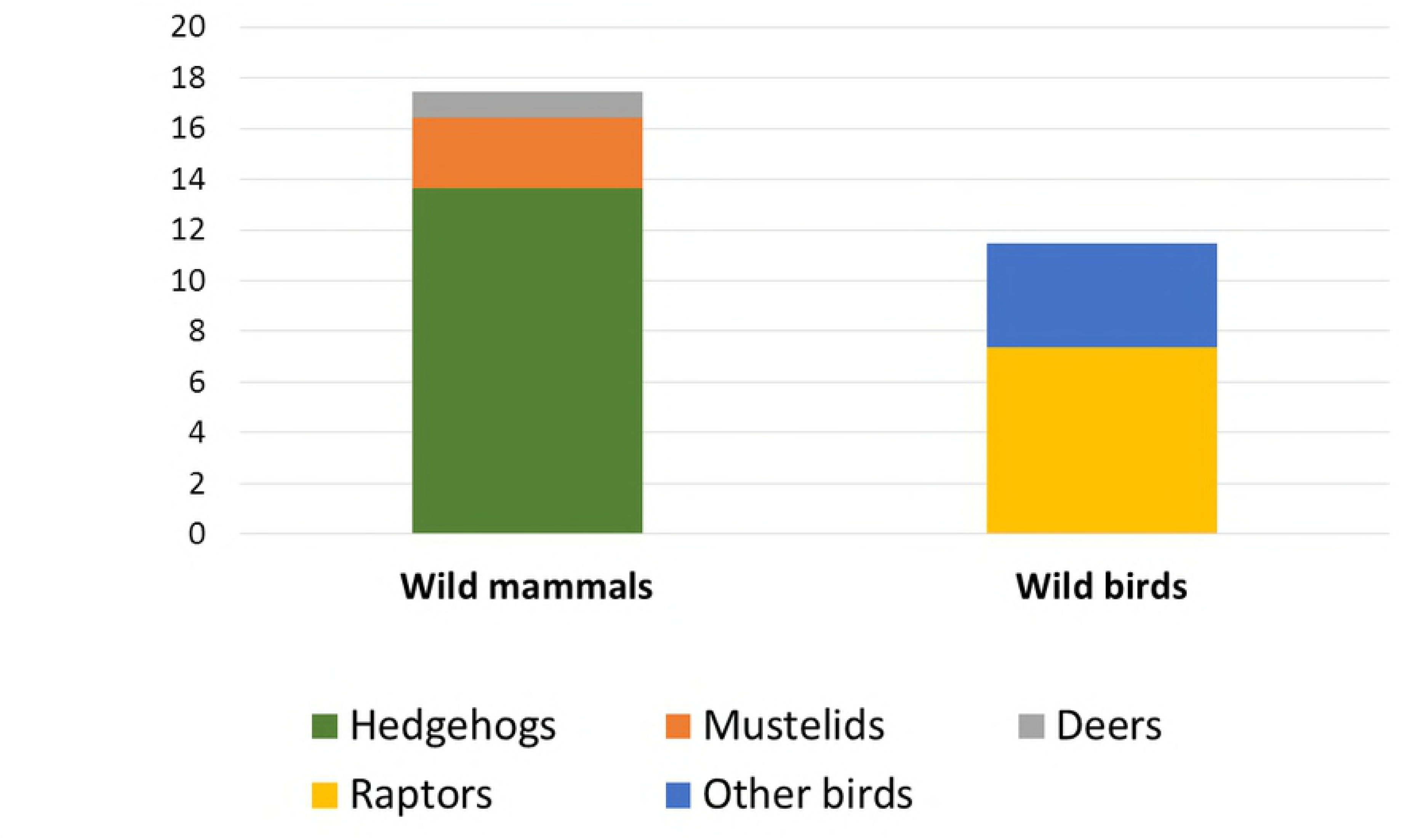
Prevalence of cephalosporin resistant (CR) bacteria in the different wildlife categories.

CR isolates belonged to several genuses within the *Enterobacteriaceae* family, with *E. coli* being detected most frequently (45%). Interestingly, other clinically relevant enterobacteria, including *K. pneumoniae* (20%), *C. freundii* (15%), *Ent. cloacae* (5%), *P. mirabilis* (5%), *Providencia* spp (5%) and *Serratia marcescens* (2.5%) were also identified as carriers of CR genes. The most common ESBL or pAmpC-encoding genes were *bla*_CMY-2_ (50% of the isolates), *bla*_SHV-12_ (23%), *bla*_CMY-1_ (20%), *bla*_TEM-1b_ (20%), and *bla*_CTX-M-15_ (18%). However, other gene variants such as *bla*_CTX-M-3,_ *bla*_SHV-1,_ *bla*_SHV-_ 11, *bla*_SHV-28_ and *bla*_SHV-167_ were also detected.

A high genetic diversity in terms of CR encoding genes was observed in all *Enterobacteriaceae* spp, with 40% (16/40) of the isolates harbouring 2 to 5 different resistance genes in the same isolate (Table 1). Furthermore, carbapenemase-encoding gene, OXA-48 was detected in *E. coli* and *P. mirabilis* isolated from European hedgehog and Barn owl, respectively (Table 1).

**Table 1.**
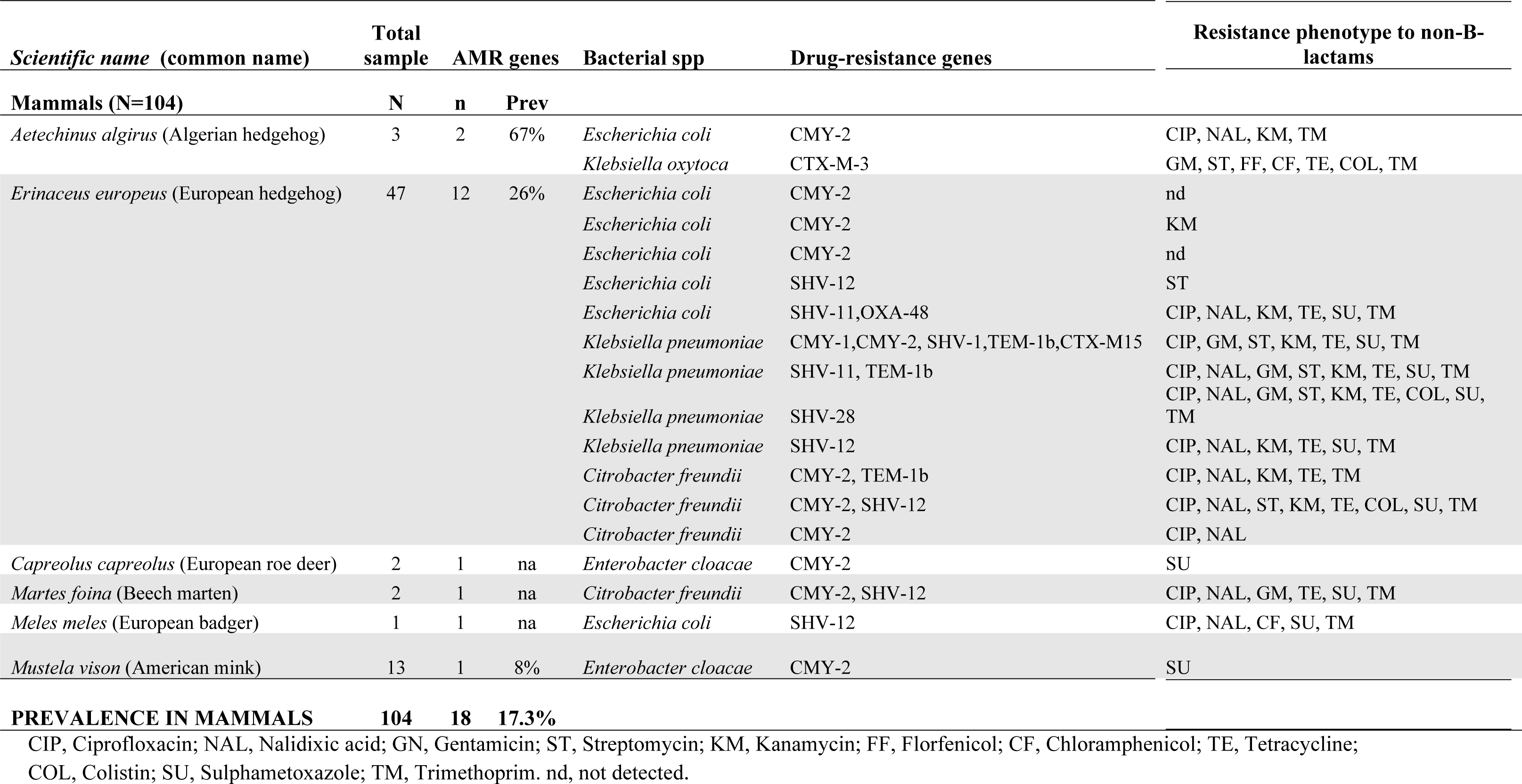
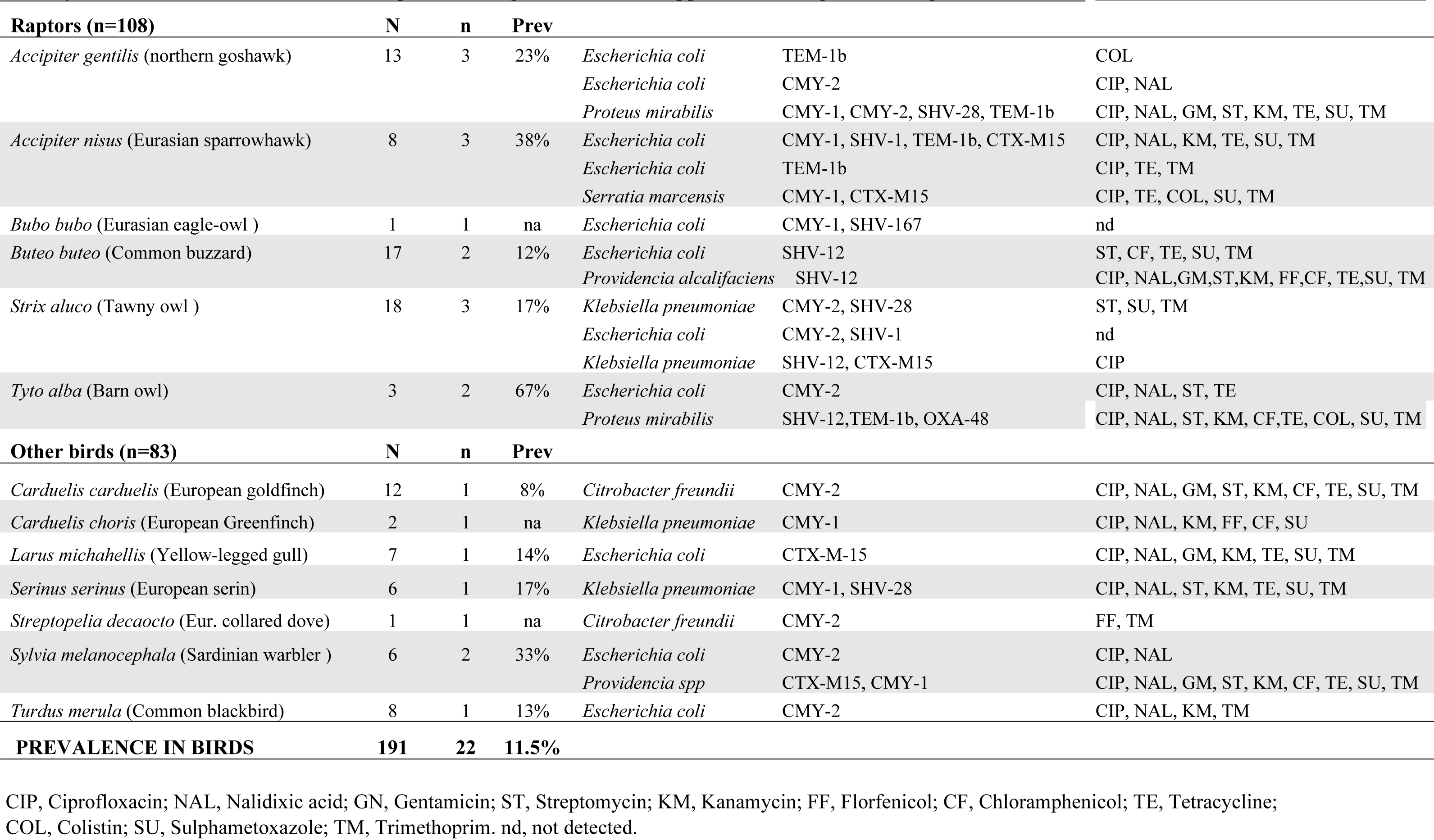
Prevalence and antimicrobial resistance genotypes and phenotypes of beta-lactamase producing *Enterobacteriaceae* spp, detected in wildlife.

Most of the ESBL/pAmpC *Enterobacteriaceae* isolates (92%), with the exception of *Ent. cloacae*, were multiresistant with a common resistance phenotype comprising β-lactams-quinolones-tetracycline-sulfamethoxazole/trimethoprim (Table 1). *K. pneumoniae* and *C. freundii* isolates both presented a multi-drug resistance profile including the resistance to aminoglycosides. Moreover, 90% of the *K. pneumoniae* isolates were resistant to ciprofloxacin and sulphametoxazole, 70% to kanamycin, 55% to streptomycin, and 10% to florfenicol. Additionally, all tested *C. freundii* isolates exhibited resistance to trimethoprim, 90% to ciprofloxacin and 80% to nalidixic acid and tetracycline (Fig 3). Although no *mcr*-1 genes were detected in this study, the colistin resistant phenotype was observed in *Klebsiella* spp isolated from a European greenfinch and Algerian hedgehog, and in a *Providencia* spp isolated from a common buzzard.

**Fig 3.**
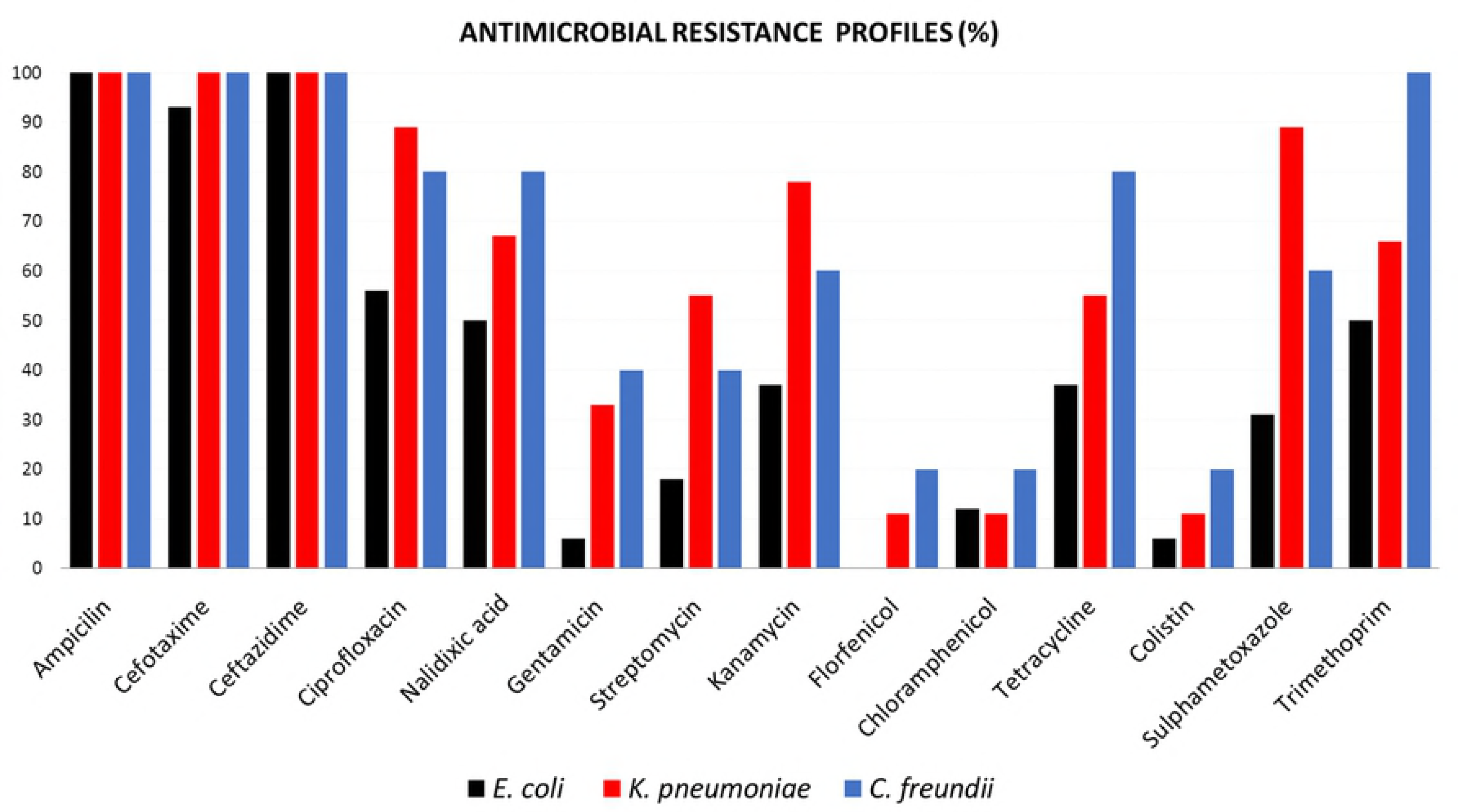
Percentage of antimicrobial resistance in ESBL producing *Enterobacteriaceae* isolates according to Minimal Inhibitory Concentration Test. Number of isolates tested: *E. coli* (n=16), *K. pneumoniae* (n=9), *C. freundii* (n=5).

## Discussion

This study identifies for the first time a high percentage of wild mammals and wild birds as carriers of different human nosocomial-like *Enterobacteriaceae* species. These isolates harboured a large diversity of ESBL/pAmpC genes, presented a high prevalence of resistance to fluoroquinolones (principally *K. pneumoniae* and *C. freundii* isolates) and in two occasions resistance to carbapenems, all drugs of last resort for the treatment of multidrug resistant infections in hospital settings [4]. Additionally, new ESBL gene variants are reported in wildlife for the first time.

*E. coli* are the most reported ESBL/AmpC-producing enterobacteria worldwide, with increasing frequency from animals, food, environmental sources and humans. In recent years, CR- *E. coli* transmission has been reported in different hosts, demonstrating a close human-animal ESBL/AmpC gene similarity between livestock (broilers and pigs) and personnel working at the farms [10]. Additionally, similar CR genes have been reported between isolates from the community and those from human clinical settings, sewage water and wild birds [10]. Although ESBL transmission has been studied extensively in *Enterobacteriaceae* from humans and livestock, data on antimicrobial resistance in the environment is still limited [2]. Moreover, most of the studies related to ESBL-carrying bacteria in wildlife are focused on the wild bird population and mainly restricted to *E. coli* species [25].

Studies performed in the Iberian Peninsula in wildlife, have reported *bla*_CTX-M-1_ as the main ESBL gene circulating [19,26]. Additionally, *bla*_CTX-M-14a_ and *bla*_SHV-12_ have also been frequently detected in *E. coli* from wild animals [12,27-29] with *bla*_CTX-M-15_ described in a recent study carried out in wild birds in Tunisia [17]. Interestingly, in the present study, *bla*_CTX-M-1_ and *bla*_CTX-M-14a_ were never detected, whereas *bla*_CMY-2,_ *bla*_SHV-_ 12 and *bla*_CTX-M-15_ were frequently isolated not only in *E. coli* but also in *K. pneumoniae* and *C. freundii* isolates. Furthermore, *bla*_CTX-M-15_ and *bla*_SHV-12_ are currently the most predominant enzymes in human clinical specimens from community and healthcare-associated infections in Spain [30,31], likely suggesting the human community as the initial source of ESBL-*Enterobacteriaceae* environmental contamination.

In this study, 6.8% of wild mammals, principally European hedgehogs and mustelids, harboured ESBL/AmpC-producing *E. coli*, the remaining 10.7% resistant isolates corresponded to other *Enterobacteriaceae* spp. Our results are in agreement with previous studies conducted in Spain reporting low to moderate (1.3%-10%) prevalence of ESBL/AmpC-producing *E. coli* genes in wild mammals [19]. In particular, in that study, hedgehogs, deer and minks were found as reservoirs of *bla*_CMY-2_ and *bla*_SHV-12_ *E. coli* variants [19,27]. However, in the present study new gene variants *bla*_CTX-M-3_, *bla*_SHV-1_, *bla*_SHV-11_, *bla*_SHV-28_ and *bla*_SHV-167_, are reported in wildlife.

Regarding the avian species analysed, the high prevalence of *bla*_SHV-12_ detected specially in raptors is in concordance with previously reported data in Spain [12]. These results confirm the successful dissemination of *bla*_SHV-12_ variants among the wild bird-population in Spain [12], The Netherlands [32], Poland [33] and the Czech Republic [34].

Plasmid-mediated colistin resistance by *mcr*-1 has been reported worldwide in *Enterobacteriaceae* isolated from humans, livestock, companion animals, food and wildlife [35]. Colistin has been used in veterinary medicine during the last decades for the treatment of gastrointestinal infections in livestock, principally in pigs and poultry [36]. Consequently, livestock is considered the main reservoir of *mcr*-1 selection and dissemination worldwide. In a recent work, whole genome sequencing based analysis disclosed the relationship among *mcr*-1-harbouring *E. coli* isolates recovered from the environment, pig production and human clinical isolates, demonstrating the rapidly evolving epidemiology of plasmid-mediated colistin-resistant *E. coli* strains worldwide and the importance of the One Health approach [37]. In our study, some *Klebsiella* and *Providencia* spp isolates were phenotypically resistant to colistin, but no *mcr*-1 gene was detected in the isolates examined. Nevertheless, although *mcr*-1 is the most commonly reported gene for colistin resistance, other less frequent genes not examined in the study, like *mcr*-2 to −5, could not be disregarded.

Information about carbapenem-resistant *Enterobacteriaceae* is very scarce in wildlife. There is a study conducted in Germany reporting a carbapenem-resistant *Salmonella enterica* from a wild bird [38]. In this study, we report the presence of *bla*_OXA-48_ in *E. coli* and *P. mirabilis* isolates from a European hedgehog and a Barn owl, respectively. Since, carbapenem-resistance genes have not yet been reported in livestock in Catalonia (these antibiotics are not authorized for animal production), the original source of these enzymes is likely to be hospitals and healthcare settings, although transmission from soil bacteria cannot be disregarded.

In this line, not many wildlife studies have reported the presence of other ESBL-producing *Enterobacteriaceae* species rather than *E. coli* in. Within them, *K. pneumoniae* has been described in low prevalence (1.5% on average) in wild gulls from different European countries [39], Chile and Canada [40], up to 23% in gulls from Alaska [41]. More recently, 8.6% wild migratory birds from Pakistan showed *bla*_CTX-M-15_ ESBL-producing *K. pneumoniae* [18]. Additionally, ESBL-producing *Enterobacteriaceae* have been described in wild birds and rodents worldwide, including ESBL-producing *K. pneumoniae* ST307 and *E. coli* ST38 clonal lineages recently reported in an urban West African rat population [42].

To our knowledge, there are no reports in wildlife describing the presence of CR genes in such a variety of *Enterobacteriaceae* spp, like *Citrobacter* spp, *Serratia* spp, or *Enterobacter* spp. *C. freundii*, is considered an opportunistic pathogen, associated with nosocomial infections, especially in patients who have been hospitalized for a prolonged period of time. In the last years, this bacterium has been classified as an emerging health care associated to urinary tract infections commonly diagnosed in healthcare settings [43]. *Ent. cloacae* has been reported as important opportunistic and multiresistant pathogen involved in outbreaks of hospital-acquired infections in Europe, particularly in France [44]. ESBL- *S. marcescens* has also been classified as one of the top ten priority pathogens causing infections in intensive care units [45]. The high prevalence of CR *Enterobacteriaceae* encountered in this study is really concerning, since wildlife is not directly exposed to any antimicrobial agents. Therefore, faecal contamination of water or soil with MDR bacteria and/or antimicrobial residues can lead to a selection pressure. Wastewaters from urban areas and hospitals have been identified as one of the major sources of AMR environmental contamination [2]. High prevalence of *bla*_SHV-12_ but also *bla*_TEM-1_ and *bla*_CTX-M-1_ alleles have been reported in aquatic environments (urban waters, natural or artificial water reservoirs, seawater or drinking water) in several countries worldwide, likely due to their relatively easy transmission to surface water through waste water treatment plant discharges [2,46]. In our study, we observed wildlife in close contact with urban and farming areas of Catalonia carrying a large variety of zoonotic/nosocomial bacteria genetically resistant to β-lactams-quinolones-tetracycline-sulfamethoxazole/trimethoprim-aminoglycosides with similar resistant genes to those found in livestock and clinical settings. Moreover, OXA-48 variants with an-extended spectrum of resistance to carbapenems were also detected in our wildlife population of Catalonia. This variant is highly prevalent in hospital settings in Spain [20].

## Conclusions

This study describes for the first time a high prevalence of *Enterobacteriaceae* spp harbouring a large variety of carbapenem and CR genes in the wildlife population of Catalonia. Bacterial spp described in this collection are associated to nosocomial infections and most of the gene-variants described here are frequently found in clinical settings. Since these wild animals had not previous antimicrobial treatment, our results suggest that both, antimicrobial residues and antimicrobial resistant bacteria are a spill-over consequence of anthropogenic pollution. Additionally, wildlife can contribute indirectly to the dissemination of resistance genes into other natural areas increasing the prevalence of AMR genes in natural environments. Thus, implementation of control measures to reduce the impact of anthropogenic pressure in the environment is urgently needed.

In summary, these results support the concept that wildlife is a good sentinel of AMR environmental contamination and simultaneously underline the importance of the One Health approach. Further studies are needed to assess clonal relatedness among different cephalosporin and carbapenem resistant enterobacteria at the human-animal-environment interface.

## Acknowledgements

Our grateful thanks to the Torreferrusa WRC staff. A. Vidal was supported by a PIF grant from the Universitat Autònoma de Barcelona. Contract of LMG was supported by the Instituto Nacional de Investigación y Tecnología Agraria y Alimentaria (INIA) and the European Social Fund.

